# Species Identification and Antimicrobial Susceptibility of Mycobacterial Isolates from Dogs and Cats

**DOI:** 10.64898/2026.01.13.699242

**Authors:** Sreekumari Rajeev, Jaclyn Azelby, Porsha Reed, Rebekah Jones, Rajeev V. Nair, Brian Johnson

## Abstract

Nontuberculous *Mycobacterium* (NTM) species are increasingly recognized as opportunistic pathogens in companion animals, yet data on species distribution and antimicrobial resistance remain limited. The goal of this study was to evaluate species identification methods and determine antimicrobial susceptibility profiles of NTM isolates from cats and dogs submitted to the University of Tennessee veterinary diagnostic laboratory. Twenty-nine NTM isolates (25 feline, 4 canine) collected between 2016 and 2022 were cultured and identified using MALDI-TOF MS and 16S rRNA sequencing. Antimicrobial susceptibility testing (AST) was performed by broth microdilution. Most isolates (27/29) were rapidly growing mycobacteria from skin or wound samples. Combined MALDI-TOF and sequencing identified 11 isolates within the *Mycobacterium fortuitum* complex, 10 as *M. smegmatis*, 4 within the *M. abscessus* group, and one each as *M. mucogenicum* and *M. farcinogenes*. MALDI-TOF failed to identify four isolates, including two slow-growing strains. AST revealed consistent susceptibility to amikacin and moxifloxacin, while resistance was highest to doxycycline (44%), clarithromycin (37%), and tobramycin (33%). Rapidly growing NTM predominates in feline cutaneous infections, with species-specific resistance patterns underscoring the need for accurate identification and AST to guide therapy.

**Importance Statement:** Nontuberculous mycobacteria (NTM) are emerging causes of skin infections in cats and dogs, yet little is known about which species are involved or how they respond to treatment. Knowing the specific species and their resistance patterns is critical for choosing effective treatments and improving outcomes for affected animals. This study provides critical baseline data by identifying NTM species and their antimicrobial resistance patterns in companion animals. These findings highlight the need for accurate diagnostic methods and targeted therapy to improve clinical outcomes. Understanding these trends is essential for guiding veterinary care and informing future research on antimicrobial resistance in animal health.

## Introduction

Non-tuberculous *Mycobacterium* (NTM) species are ubiquitous environmental organisms increasingly recognized as opportunistic pathogens in both humans and animals[1, 2]. In veterinary medicine, NTM infections are most frequently reported in cats and dogs, typically associated with cutaneous and subcutaneous lesions, draining tracts, and occasional respiratory involvement[3–7]. These infections often follow local penetrating trauma-related infections or surgical wound infection procedures, allowing inoculation of organisms from contamination from soil or water into tissues[1]. There are several reports of infection with members of *Mycobacterium tuberculosis* complex, most commonly *M. bovis,* in cats[8, 9]. In contrast, NTM infections pose diagnostic and therapeutic challenges due to their diverse species distribution and variable antimicrobial susceptibility profiles[1]. Rapidly growing mycobacteria (RGM), including members of the *M. fortuitum* complex, *M. smegmatis*, and *M. abscessus* group, are commonly implicated in feline and canine infections[4]. These organisms exhibit intrinsic resistance to many antibiotics and unpredictable susceptibility patterns, making empirical therapy unreliable[1]. Accurate species identification and antimicrobial susceptibility testing (AST) are therefore essential for guiding treatment. However, identification of NTM in veterinary diagnostic laboratories remains challenging because of the need for specialized methods[2]. Traditional phenotypic approaches are time-consuming and often insufficient for species-level resolution. Matrix-Assisted Laser Desorption/Ionization Time-of-Flight Mass Spectrometry (MALDI-TOF MS) has emerged as a rapid and cost-effective tool for bacterial identification, including NTM, but its performance depends on extraction protocols and database coverage[10, 11]. Molecular methods such as 16S rRNA sequencing remain the gold standard for definitive identification, particularly for slow-growing species or isolates with ambiguous MALDI profiles. In parallel, standardized AST methods, such as broth microdilution interpreted according to Clinical and Laboratory Standards Institute (CLSI) guidelines, provide critical data for therapeutic decision-making. Despite these advances, published data on NTM infections in companion animals, particularly species distribution and antimicrobial resistance patterns, remain limited, other than a few case-based or region-specific reports and few studies have systematically evaluated diagnostic workflows combining MALDI-TOF and sequencing with AST in veterinary settings. In this study, we characterized NTM isolates from cats and dogs submitted to a university veterinary diagnostic laboratory and evaluated the performance of MALDI-TOF MS and targeted sequencing for species identification and determined antimicrobial susceptibility profiles using standardized methods.

## Materials and Methods

### Bacterial Isolates

We analyzed 29 nontuberculous *Mycobacterium* spp. isolates obtained from dogs and cats submitted to the University of Tennessee Veterinary Diagnostic Laboratory between 2016 and 2022. Isolates included both in-house and referral cases. Frozen stocks were subcultured on 5% sheep blood agar and incubated aerobically at 37 °C with 5% CO₂ for up to 7 days. Colonies were examined using Gram stain and Kinyoun modified acid-fast stain. Patient information associated with each isolate was retrieved from laboratory records.

### Species Identification

Species identification was performed using Matrix-Assisted Laser Desorption/Ionization Time-of-Flight Mass Spectrometry (MALDI-TOF MS) with the MycoEx extraction method as recommended by the manufacturer (Bruker, Billerica, MA), omitting the heat inactivation step. Briefly, colonies were washed twice with 300 μL HPLC-grade water and 900 μL ethanol, then centrifuged at 13,000 rpm for 2 min. The pellet was air-dried for 3 min, mixed with zirconia/silica beads, and treated with 50 μL acetonitrile by vortexing for 1 min. After adding 50 μL of 70% formic acid, samples were centrifuged at 13,000 rpm for 2 min. One microliter of supernatant was spotted onto a MALDI target plate and analyzed using a MALDI Biotyper system.

DNA was extracted using the Quick-DNA Fungi/Bacteria Miniprep Kit (Zymo Research) following the manufacturer’s protocol. PCR amplification was performed using two sets of previously described 16s rRNA-based primer sets [(MB1-5’- AGT GGC GAA CGG GTA AGT AAC-3’/MB7- 5’ TTA CGC CCA GTA ATT CCG GAC AA-3’) and (MB246-5-’AGA GTT TGA TCC TGG CTC AG-3’-/MB247-5’-TTT CAC GAA CAA CGC GAC AA-3’)][12]. PCR amplification using MB1/MB7 and MB246/MB247 primer sets yields 469 and 590-basepair amplicons, respectively. Amplicons were treated with ExoSAP-IT (Thermo Fisher Scientific) and sequenced via Sanger sequencing by a commercial service. Resulting sequences were queried against the NCBI database using BLAST.

### Antimicrobial Susceptibility Testing

Broth microdilution AST was performed using Sensititre RAPMYCO plates (Thermo Fisher Scientific) for 27 rapidly growing isolates against 12 antimicrobial agents; testing was not performed on two slow-growing isolates. Colonies were suspended in sterile water and adjusted to a 0.5 McFarland standard using a nephelometer. Fifty microliters of suspension were added to cation-adjusted Mueller-Hinton broth with TES buffer to achieve ∼5 × 10⁵ CFU/mL. Aliquots (100 μL) were dispensed into each well, plates were sealed and incubated at 30 °C in ambient air for ≥72 h, and results were read using the BIOMIC V3 system (Giles Scientific, Santa Barbara, CA).

## Results

### Patient Information and Source of Isolates

A total of 29 nontuberculous *Mycobacterium* isolates were analyzed, comprising 25 from cats and 4 from dogs. The animals ranged in age from 6 months to 14 years. Sample sources included skin (n = 21), wounds (n = 4), lymph node (n = 1), and tracheal wash (n = 1). The most common presenting complaints were non-healing wounds or skin lesions (n = 23) and chronic upper respiratory infections (n = 2); four cases lacked clinical details. Colonies appeared within 72 hours for 27 isolates, which were classified as rapid-growing mycobacteria, while two isolates required 7 days for visible growth and were designated as slow-growing. All isolates were Gram-positive and acid-fast positive (9 complete, 20 partial).

### Species Identification

MALDI-TOF MS identified 25 isolates to species level; four isolates, including the two slow-growing strains, were not identified by MALDI but were confirmed by sequencing. Primer set 2 demonstrated better concordance with MALDI results compared to primer set 1. Combined MALDI-TOF and 16S rRNA sequencing identified 11 isolates within the *Mycobacterium fortuitum* complex, 10 as *M. smegmatis*, 4 within the *M. abscessus* group, 1 as *M. mucogenicum*, and 1 as *M. farcinogenes*. Nine isolates matched across all three identification methods, and one additional isolate matched by both primer sets. Full species identification data are summarized in Detailed isolate descriptions, origins, and MALDI-TOF identification scores are provided in **Supplemental file 1**.

### Antimicrobial Susceptibility Testing

AST results were interpreted using CLSI M24 guidelines. All *M. smegmatis* isolates (n = 10) were susceptible to amikacin, imipenem, moxifloxacin, and trimethoprim-sulfamethoxazole, with the highest resistance observed to clarithromycin. Isolates within the *M. fortuitum* complex (n = 11) were uniformly susceptible to amikacin and moxifloxacin, with resistance observed primarily to tobramycin and doxycycline. The *M. abscessus* group (n = 4) exhibited multidrug resistance, including resistance to moxifloxacin and doxycycline. The isolates identified as *M. mucogenicum* and *M. farcinogenes* were resistant to doxycycline. Overall, the highest resistance was observed to doxycycline (44%), followed by clarithromycin (37%) and tobramycin (33%). Figure 1 represents the overall antimicrobial susceptibility patterns of all the isolates tested. Figure 2 represents the summary of antimicrobial susceptibility patterns of isolates characterized into species groups, and the complete data with all MIC values are shown in the supplemental table 2.

**Figure 1:**
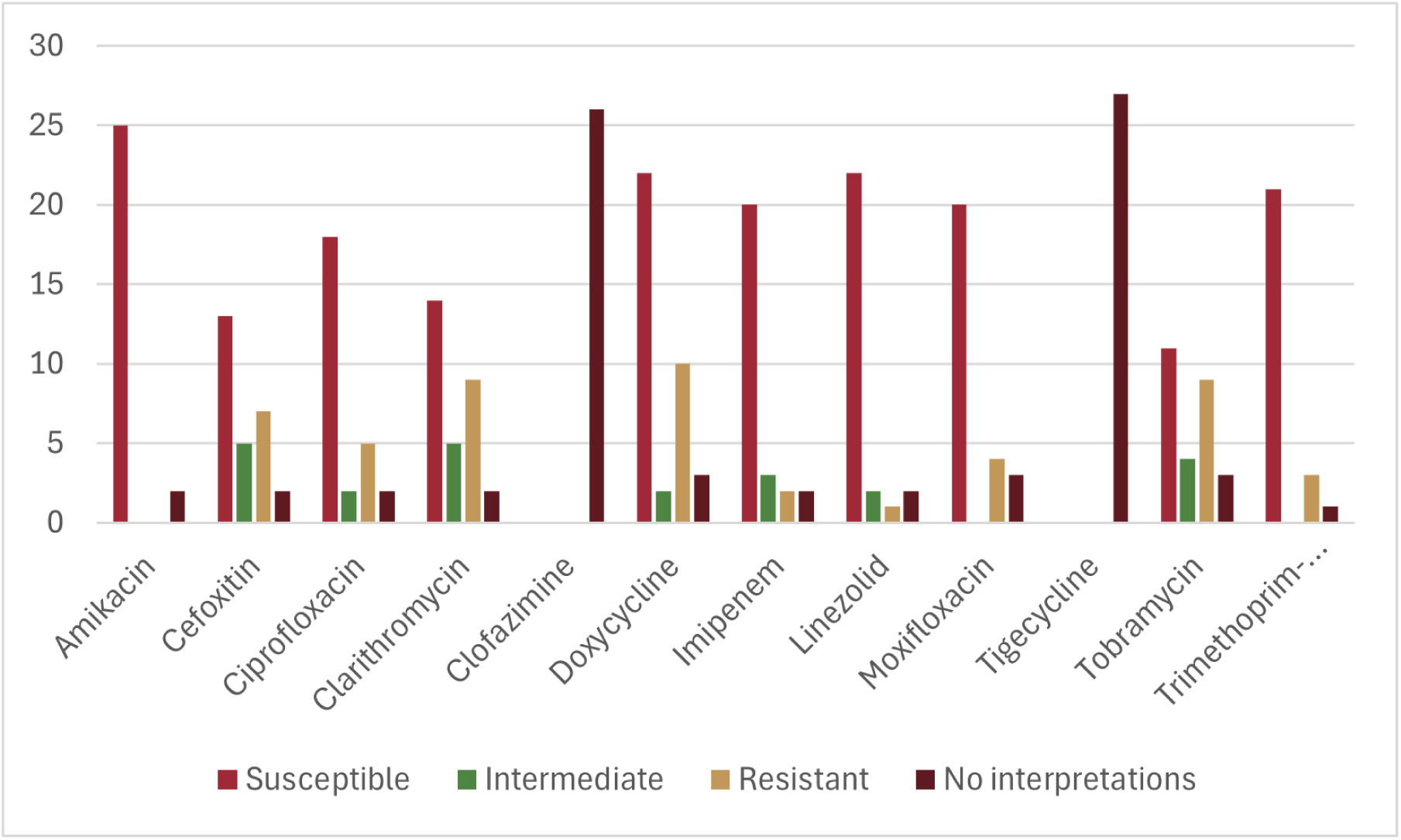
The overall antimicrobial susceptibility patterns of all the isolates tested.

**Figure 2:**
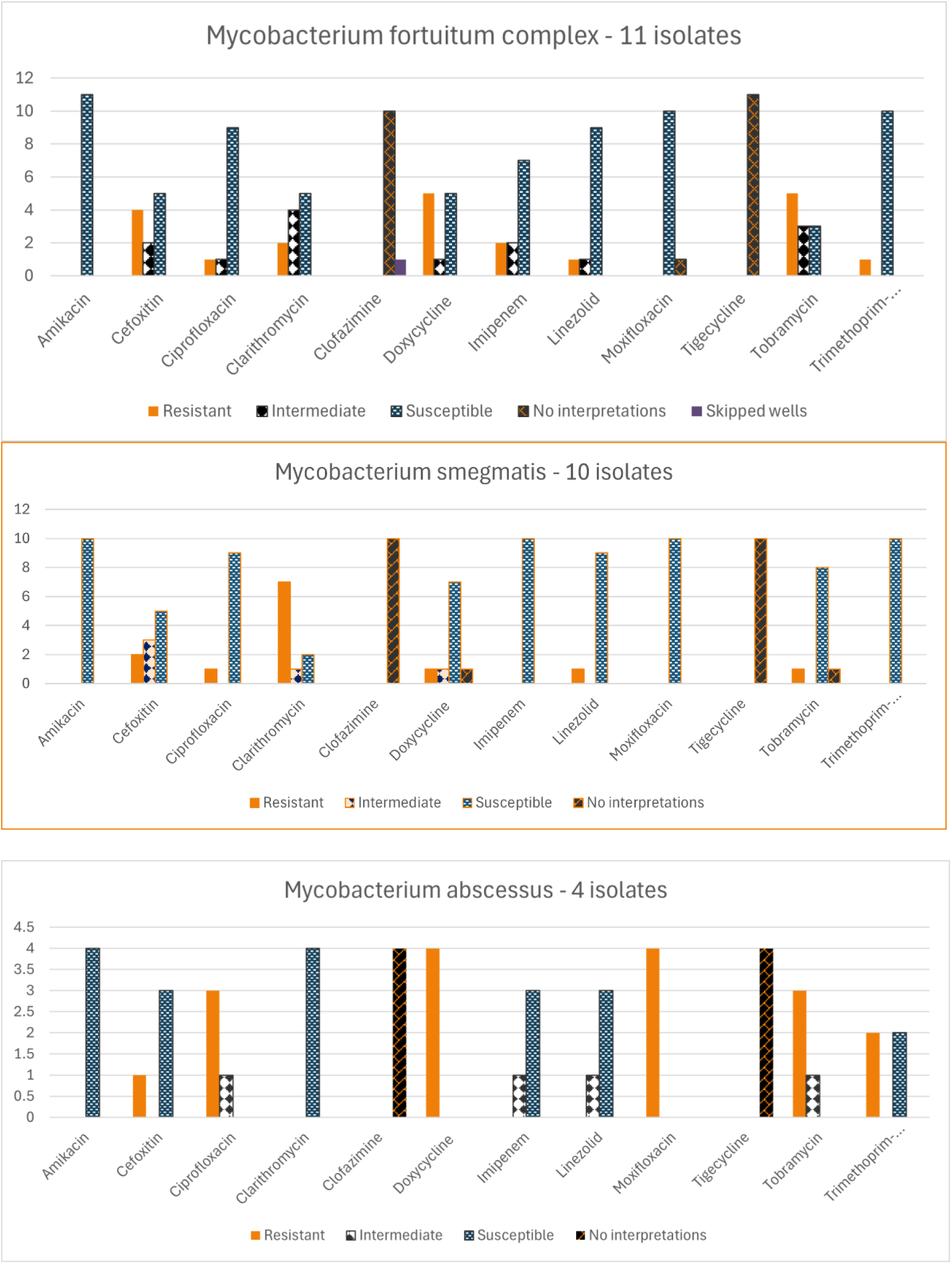
The summary of antimicrobial susceptibility patterns of isolates categorized into major species groups.

## Discussion

Our workflow combined MALDI-TOF MS with targeted sequencing, reflecting a pragmatic approach in veterinary diagnostics where rapid turnaround must be balanced against the known limitations of MALDI for certain mycobacterial taxa. In our study, MALDI failed to identify four isolates, notably including the two slow-growing strains, whereas sequencing resolved species-level identities consistent with evidence that MALDI-TOF demonstrates higher accuracy for RGM than for slow-growing NTM, and that library coverage critically influence performance. Multicenter evaluations and best-practice guidance concur that while MALDI-TOF provides accurate, reproducible genus/species identifications for most clinically relevant NTM, closely related or underrepresented species may remain problematic, warranting confirmatory molecular testing. Continued improvements in commercial libraries and modules (e.g., Bruker’s dedicated mycobacteria library and workflows) are expanding coverage, but laboratories should maintain dual-method strategies when identification impacts treatment decisions. Concordance patterns in our data (better agreement with primer set 2) emphasize that target choice matters for NTM speciation; the literature similarly notes that single-gene targets (e.g., 16S) may be insufficient for certain taxa, and secondary loci (e.g., *rpoB*) can resolve ambiguities. These findings support a tiered algorithm for MALDI-TOF for rapid provisional ID, followed by sequence confirmation for slow-growing isolates, atypical spectra, or species with therapeutic ramifications.

The predominance of *M. fortuitum* complex and *M. smegmatis* among feline skin/wound isolates is consistent with veterinary reports that these RGM are frequent cutaneous opportunists in cats and dogs. The identification of a smaller *M. abscessus* subset is notable because this group is intrinsically more drug-resistant and poses greater management challenges; its emergence in companion animals mirrors human medicine, where *M. abscessus* has become a major NTM pathogen. Although rare in our series, isolates of *M. mucogenicum* and *M. farcinogenes* reinforce the taxonomic breadth of NTM encountered in veterinary patients and underscore the need for species-level reporting to guide therapy. AST results, interpreted per CLSI M24/M24S, demonstrated reliable activity of amikacin across major RGM groups and consistent susceptibility to moxifloxacin in *M. fortuitum* complex findings that align with established performance standards and published susceptibility summaries for RGM. In contrast, we observed high resistance to doxycycline (overall 44%) and substantial resistance to clarithromycin (37%) and tobramycin (33%). Doxycycline’s limited clinical utility against RGM is well recognized and reflects class-wide limitations of tetracyclines for many NTM, supporting avoidance of empiric doxycycline pending AST. In our study, we used the standard 3-day incubation period for AST. To assess the inducible macrolide resistance mediated by the *erm* (41) gene, a 7-day read is recommended. [13]. The shorter incubation period used might have potentially led to false susceptibility. Molecular characterization of erm determinants in these cases is increasingly recommended to complement phenotypic AST, given the complex resistance mechanisms and their impact on treatment strategies based on the AST results[14]. In companion animals, recent regional studies similarly document variable macrolide activity in *M. fortuitum* complex and multidrug resistance in *M. abscessus*, reinforcing the need for species-and subspecies-level ID plus resistance gene detection where feasible[4]. Our use of standardized commercial plates and BIOMIC V3 for broth microdilution reading/interpretation provides standardized CLSI-based calls and an image-backed audit trail, which is advantageous for quality assurance and longitudinal AMR surveillance. Nonetheless, laboratories should remain vigilant for inducible phenotypes (e.g., macrolides in *M. abscessus*) and ensure incubation durations and AST read schedules are adequate for organism-specific behavior. Other limitations of this study include a modest sample size, a focus on a single geographic region, and the absence of clinical outcome data to correlate AST with therapeutic success, restrictions common to veterinary NTM studies, and emphasizing the need for multicenter datasets. Additionally, while MALDI coverage is improving, library gaps and closely related taxa can still produce discordant identifications; our discrepant cases highlight the value of confirmatory sequencing. Future work should expand regional surveillance of RGM in companion animals to define species distribution and resistance trends across settings, incorporate molecular resistance profiling into routine workflows for *M. abscessus* and other problematic taxa, and evaluate clinical outcomes relative to species-level ID and AST-guided therapy regimens. The standardization of MALDI protocols and adoption of enhanced libraries will likely improve identifications, while targeted sequencing panels can resolve ambiguity, together optimizing turnaround time in veterinary diagnostic practice. Whole genome sequencing (WGS) is increasingly becoming the standard for bacterial species identification due to its superior resolution compared to 16S rRNA sequencing. While 16S sequencing targets a single conserved gene, making it cost-effective and suitable for genus-level identification or microbiome profiling, it often fails to differentiate closely related species or strains. In contrast, WGS not only enables precise species and strain-level taxonomy but also provides critical insights into antimicrobial resistance and virulence factors. Although WGS is more expensive and computationally demanding, its ability to deliver comprehensive genomic information makes it indispensable for clinical diagnostics, outbreak investigations, and evolutionary studies. Therefore, subjecting our isolates to WGS is imperative for accurate identification and functional characterization.

## Acknowledgments

We would like to thank UTCVM, the Center of Excellence in Livestock Diseases & Human Health, for supporting the project.

